# Vector Semantics of Multidomain Protein Architectures

**DOI:** 10.1101/2025.07.07.663606

**Authors:** Xiaoyue Cui, Yuting Xiao, Maureen Stolzer, Dannie Durand

## Abstract

Multidomain proteins are mosaics of *domains*, protein modules that are associated with a specific structure or function and are found in diverse combinations. This modular organization facilitates the evolution of novel protein functions, but the principles that govern the relationship between the domain content of a protein and its function is poorly understood. In particular, do domains always contribute the same function, or does the functional contribution of a domain depend on the neighboring domains in the protein? To answer this question, we used vector embeddings, which account for local contextual signals, to model the protein domain content of multidomain proteins. We observe that multidomain architectures that are semantically similar share more functional attributes than multidomain architectures selected based on domain content similarity, alone, suggesting that context is important for understanding the relationship between domain content and protein function. Surprisingly, vector semantics also identified multidomain architecture pairs with significantly high functional similarity, despite having no domains in common at all, suggesting that vector semantics may be discovering domain “synonyms”. Taken together, our results underscore the importance of contextual models for understanding the interplay between domain architecture evolution and functional innovation in multidomain proteins.

## Introduction

Multidomain proteins are mosaics of structural or functional units called domains. Domains act as protein modules, are found in otherwise unrelated proteins, and fold independently in many different contexts. The *domain architecture* (DA) of a protein is an abstract representation consisting of its domains in N- to C-terminal order. For example, the domain architecture of the Mpp6 protein shown in Fig. 1 can be represented as `L27-L27-PDZ-SH3-GuK`. The domain architecture abstraction is widely used to probe questions of protein evolution, including variation in the domain repertoire across taxonomic lineages [Karev et al., 2004, Tordai et al., 2005, Ye and Godzik, 2004, Cromar et al., 2016, Dohmen et al., 2020], plasticity in domain order [Bashton and Chothia, 2002, Kummerfeld and Teichmann, 2009, Weiner 3rd et al., 2006], domain occurrence graphs [Vogel et al., 2005, Karev et al., 2002, Cromar et al., 2014, Przytycka et al., 2006], and domain promiscuity, i.e., the propensity of a domain to co-occur with many other domains [Marcotte et al., 1999, Basu et al., 2009, 2008, Cohen-Gihon et al., 2011, Cromar et al., 2014, Weiner et al., 2008].

**Fig. 1.**
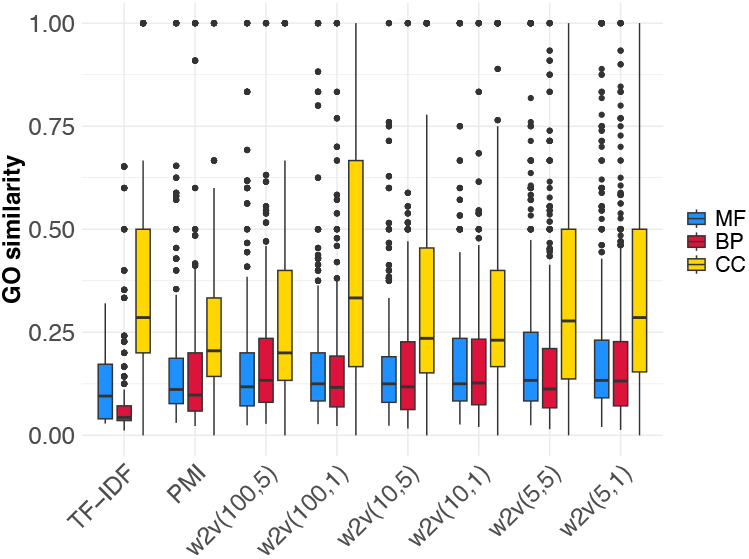
Functional similarity between target domain architectures and the *k* = 1 nearest neighbor that lacks a common domain.

Multidomain sequences evolve via the insertion, internal duplication, and deletion of domains, resulting in families with similar domain architectures, but with some variation in domain content, order and copy number (e.g., the Membrane Associated Guanylate Kinases (MAGuKs) [Te Velthuis et al., 2007] shown in Fig. 1).

The modular organization of multidomain proteins supports rapid evolutionary exploration of new protein functions through the formation of different domain combinations that enable diverse biological roles. At the same time, this exploration is not unconstrained: the number of domain combinations observed in the multidomain repertoire is much smaller than expected by chance [Yu et al., 2019, Cui et al., 2022, Vogel et al., 2005]. The principles that govern which domain combinations are realized in nature are poorly understood. Many forces could be contributing to those constraints, including structural compatibility, mutational processes, and cellular environment.

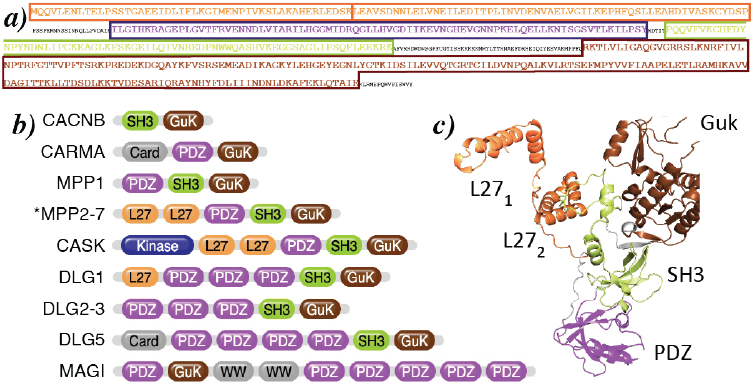
Domain architectures of MAGuK subfamilies exhibit similar domain content in various combinations. (c) Structure of MPP6, with individual domains color coded as in (b).

Here we focus on the relationship between the domain content of a protein and its function. In specific cases, the function of multidomain proteins can be understood in terms of the functions of individual domains. For example, the extracellular domains of a receptor kinase recognize a molecular signal; the intercellular kinase domain passes on that signal by phosphorylating an amino acid. More generally, proteins with identical domain architectures have more similar GO annotations, on average, than those that do not and proteins that share at least one domain have greater GO similarity than those that have no domains in common [Radivojac, 2022], suggesting that information is there to exploit. This is further supported by recent successes with protein function prediction methods that incorporate domain content information [e.g. You et al., 2018, Ibtehaz et al., 2023, Talo and Bozdag, 2025, Wang et al., 2025, Hallee et al., 2024, Chen and Luo, 2024, Hu and Zhao, 2025]. However, it is not clear whether a given domain confers the same functional properties in all contexts, or whether the functional contribution of a domain varies depending on neighboring domains in the protein.

### Vector Semantics

Here we investigate *vector semantic* models, developed for natural language processing (NLP), as an analytical framework for probing the relationship between the domain content and the function of a multidomain protein. Just as documents are strings of words, domain architectures are strings of domains, suggesting that NLP models are a promising approach for exploratory analysis of the multidomain universe. Vector semantics are based on the assumption that the meanings of words are implicitly encoded in their usage in the language. The distribution of words across documents is a representation of word usage; hence, words that occur in similar contexts tend to have related meanings. Contextual information can be encoded as a *word embedding*, where each word corresponds to a vector that represents its distribution in a corpus of documents. The goal is to select a vectorization such that words that are proximal in the vector space will tend to have related meanings. This includes words with similar meanings (e.g. chilly, cool), but also words that have meanings that are dissimilar, but are topically related (e.g., *bus, train*; *dog, animal*; or *cup, coffee*).

Embeddings used in information retrieval are sparse, high dimensional encodings. In a term frequency - inverse document frequency (TF-IDF) encoding, each dimension corresponds to a document; the *i*^*th*^ component of a word vector represents its frequency in document *i*, normalized by its frequency across the corpus [Luhn, 1957, Sparck Jones, 1972]. In a TF-IDF vectorization, words that occur in the same documents will be close in the vector space. In a Positive Pointwise Mutual Information (PPMI) [Church and Hanks, 1990] embedding, each dimension corresponds to a word in the vocabulary; the *i*^*th*^ component of a word vector is a function of its frequency of co-occurrence with the *i*^*th*^ word in the vocabulary, relative to the independent overall frequencies of the two words. In a PPMI embedding, words will be proximal if they co-occur with the same words. Static language embeddings, e.g. Word2Vec [Mikolov et al., 2013], learns compact vector representations from training data (e.g., document corpora) and more fine-grained contextual information is exploited.

Given a word embedding, a *document embedding* can be constructed, in which each document is represented as a vector obtained by combining the vectors of the words that appear in the document. Here, the goal is a vectorization such that documents on similar subjects are proximal. The words used in proximal documents will tend to have meanings that are related. Document embeddings provide a framework for comparing documents and investigating the structure of a corpus.

Multidomain proteins poses challenges for bioinformatic techniques that use sequence analysis to extract functional, structural, and evolutionary information. In families with variable domain architectures, such as the MAGuKs, some sequence segments are homologous and have discernible sequence similarity; other regions are not alignable. As a result, bioinformatic tools based on sequence comparison cannot be directly applied to multidomain protein families with mosaic sequences. At the same time, the variations in domain composition carry information that current tools are not designed to exploit. Examining word context using vector semantics provides information about the relationships between words, and hence documents, that is independent of whether the words contain similar letters or are derived from the same root in an ancient language, suggesting the promise of this approach for multidomain protein architectures

Domain embeddings [Buchan and Jones, 2020, Melidis and Nejdl, 2021] have been used to investigate the potential of vector semantics for transferring functional information between neighboring domains. Moreover, domain embeddings have been shown to improve functional prediction of proteins [Ibtehaz et al., 2023, Wang et al., 2025]. For instance, Ibtehaz et al. [2023] learned domain embeddings from domain-GO co-occurrence data, achieving state-of-the-art results on GO annotation tasks. Further integrating complementary features such as sequence similarity and protein interaction information enhanced prediction accuracy. These results support the view that domain composition carries meaningful functional signals.

### Our contributions

Here we investigate the use of domain architecture embeddings for studying functional relationships between multidomain proteins, using human domain architectures as a case study. The human genome is studied extensively and has high quality gene and domain annotations. In addition, there are many sources of detailed information about human multidomain protein families from which benchmarks can be constructed. Restricting the study to a single genome results in a relatively small corpus. However, data sets that span multiple genomes have biases that are hard to characterize and correct. It is difficult to determine whether the same domain architecture, found in multiple related genomes, arose through convergent formation of that architecture in independent lineages or via vertical descent from the same domain architecture in a common ancestor. In the latter case, the multiple instances of the domain architecture represent observations of the same event and should be discarded. Uneven taxon sampling in the underlying database further exacerbates this problem. In addition, if the functional roles of the domains change over the course of evolution, combining information from distant genomes would further confound the signal.

While the analogy between documents as strings of words and domain architectures are strings of domains is compelling, the characteristic scales of domain architecture data and natural language corpora are very different. In natural language corpora, sentences are typically 15 to 20 words in length, taken from a vocabulary of hundreds of thousands of words. Depending on the application, the document data available is almost unlimited. In contrast, the average length of a human domain architecture is less than 5 (median = 3), drawn from ∼1100 domains. There are approximately 5000 unique domain architectures in the human genome. With this in mind, we experimented with a number of domain embedding strategies, with modifications tuned to the scale of our data. We used two sparse embeddings, TF-IDF and Pointwise Mutual Information (PMI). For natural language applications, pointwise mutual information is typically restricted to positive values, focusing on words that co-occur more often than expected. In domain architectures, underrepresentation of domain pairs can be informative. Further, problems with underflow that occur with low frequency word pairs do not arise because the small scale of DA data. We also experimented with Word2Vec using a skip-gram model, which accounts for local context; i.e., the domain order and content in a window of width 2*w* + 1 centered on the current domain. In addition to the default window size of *w* = 5, we experimented with a smaller window (*w* = 1) to allow for the large number of domain architectures of length 5 or shorter. To account for the reduced scale of domain architecture data, we considered smaller dimensionalities (5 and 10), as well as the default value (100). We did not consider contextual embeddings (e.g, BERT representations, Devlin et al. [2018]), which account for contextual differences in meaning (e.g., homonyms) because most domain architectures are too short to benefit from that level of representational learning.

We apply this vector semantic framework to ask three questions about the relationship between domain content and protein function. First, do proteins with similar domain content also perform similar functions? To assess the association between domain architecture and protein function, for each embedding, we asked how accurately the function of a multidomain protein is predicted by the GO annotations of its neighbors in the embedding. Next, we asked whether this association is stronger when domain content similarity is assessed using vector embeddings, which carry implicit contextual information, compared with a direct comparison of protein domain content using Jaccard similarity, which provides a measure of the domains shared by two architectures, but does not use any of the additional information captured by a vector semantic embedding.

Finally, we considered the case where neighboring domain architectures have no domains in common. In the natural language analogy, embeddings that capture words with similar or related meanings can address the “vocabulary problem”, where an information retrieval request fails because the word usage in the query does not match the word usage in the desired document [Furnas et al., 1987]. We ask whether an analogous “domain vocabulary problem” exists for multidomain protein function and, if so, whether domain architecture embeddings can help to solve it.

## Materials and methods

### Dataset and pre-processing

Domain annotations (start and end positions) for 58, 023 ENSEMBL protein sequences in the human genome (assembly GRCh38.p13) were downloaded from the SUPERFAMILY database, version 1.75 [Pandurangan et al., 2019]. SUPERFAMILY uses a hierarchical classification based on SCOP, in which domains are grouped into families and superfamilies. Domains within the same superfamily share a structural core, although they may have low sequence similarity. In this work, we used the superfamily-level assignments. The pre-processing module of DomArchov [Cui et al., 2022] was used to extract the domain architectures of these sequences, resulting in a set of *N* = 5,031 unique DAs, with an average length of 4.2 domains per DA. These architectures comprise a set of 1,109 distinct domain superfamilies, denoted *D*.

GO terms associated with the ENSEMBL protein sequences were obtained from ENSEMBL [Martin et al., 2023] and assigned to the corresponding domain architectures. Most DAs correspond to multiple ENSEMBL sequences. In these cases, the DA is annotated with those GO codes that are associated with at least 50% of all corresponding protein sequences. The list of GO associations is expanded to include to ancestors in the GO hierarchy connected by “is a” and “part of” relations. This resulted in 1,157 distinct Molecular Function (MF), 1,726 Biological Process (BP), and 378 Cellular Component (CC) GO codes are assigned to human DAs. Of 5,031 DAs, 4,244 have at least one GO code. On average, each DA is mapped to 21 GO terms. Within the individual ontologies, 3,910 DAs have at least one MF term; 2,631 DAs have at least one BP term; 1,295 DAs have at least one CC term. Any DA with only a protein-binding (GO:0005515) annotation is removed from analyses for the MF ontology following the practice discussed in Zhou et al. [2019].

### Domain Embedding

Domain embeddings were constructed from the 5,031 unique DAs using eight variants of three different vectorization strategies, where each domain *D*_*i*_ ∈ *D* is represented as a vector, ***e***(*D*_*i*_) = [*e*_1_(*D*_*i*_), …, *e*_*m*_(*D*_*i*_)]. The dimensionality, *m*, depends on how the embedding is constructed.

### Pointwise mutual information (PMI)

The pointwise mutual information [Church and Hanks, 1990] of domain *D*_*i*_ followed by *D*_*j*_ is defined to be

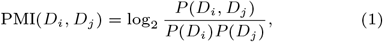

where *P* (*D*_*i*_) is the empirical frequency of domain superfamily *D*_*i*_, and *P* (*D*_*i*_, *D*_*j*_) is the empirical frequency of the bigram *D*_*i*_*D*_*j*_. A pseudocount *ψ* is added to the count of all bigrams. Following Cui et al. [2022], we use *ψ* = 0.0009.

In the PMI embedding, the *j*^*th*^ element of ***e***(*D*_*i*_) is *e*_*j*_ (*D*_*i*_) = PMI(*D*_*i*_, *D*_*j*_). The dimensionality of this embedding is the number of unique domain superfamilies in the training data, *m* = |*D*|.

### Term frequency-inverse document frequency (TF-IDF)

The inverse document frequency [Sparck Jones, 1972] of *D*_*i*_ is defined as idf(*D*_*i*_) = *N/*df(*D*_*i*_), where df(*D*_*i*_) is the number of DAs in which *D*_*i*_ occurs and *N* is the total number of domain architectures in the dataset. For any domain *D*_*i*_ and domain architecture *A*_*j*_, the term frequency tf(*D*_*i*_, *A*_*j*_) is the number of copies of *D*_*i*_ in architecture *A*_*j*_ [Luhn, 1957]. Here the *j*^*th*^ element of ***e***(*D*_*i*_) is *e*_*j*_ (*D*_*i*_) = tf(*D*_*i*_, *A*_*j*_) · idf(*D*_*i*_) and *m* = *N*.

### Word2vec

Word2Vec domain embeddings were constructed using the Python library gensim [Řehůřek and Sojka, 2010] with the skip-gram model [Mikolov et al., 2013] and a minimum word count of 1. Embeddings were calculated for vector sizes of *m* ∈ {5, 10, 100} and window sizes of *w* ∈ {1, 5}, resulting in six different Word2Vec embeddings. Each model is denoted by w2v(*m, w*). The models were trained for 15 epochs.

### Domain architecture embedding

A domain architecture embedding was constructed for each of the eight domain embeddings (i.e, TF-IDF, PMI, and 6 Word2Vec models) described above. Given domain architecture *A* = *D*_1_ … *D*_*n*_, the embedding vector of *A* is obtained by averaging the embeddings of its constituent domains

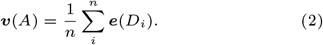

### Assessment

For assessments of functional similarity, only annotated multidomain architectures (i.e., DAs with 2 or more domains and at least one GO term) were considered. The numbers of annotated multidomain architectures in each of the three sub-ontologies are 3,193 (MF), 2,066 (BP), and 1,001 (CC).

Let *A*_*O*_,*O* ∈ {MF, BP, CC} denote the set of annotated multidomain architectures in ontology *O*. Assessment centers around the functional attributes of the *k* nearest neighbors of a target domain architecture *A* ∈ *A*_*O*_,*O* ∈ {MF, BP, CC}.

### Pairwise domain content similarity

Given a pair of domain architectures, *A*_*i*_ and *A*_*j*_, proximity in the embedding space is quantified using pairwise cosine similarity, denoted by *S*_*C*_ (***v***(*A*_*i*_), ***v***(*A*_*j*_)).

The similarity of two architectures can also be quantified based on shared domain content, independent of an embedding, using the Jaccard similarity:

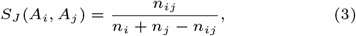

where *n*_*ij*_ is the number of domain copies shared by *A*_*i*_ and *A*_*j*_. If *A*_*i*_ and *A*_*j*_ share no domains, then *S*_*J*_ (*A*_*i*_, *A*_*j*_) = 0.

### Functional similarity

#### Comparison of two domain architectures

For each ontology, let ℱ_*O*_(*A*) denote the set of GO terms annotated to domain architecture *A*. The functional similarity between domain architectures *A*_*i*_ and *A*_*j*_, is defined to be the Jaccard similarity between ℱ_*O*_(*A*_*i*_) and ℱ_*O*_(*A*_*j*_):

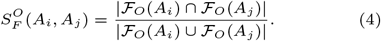

#### Comparison with sets of domain architectures

The GO term annotation of a set domain architectures 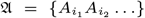 is defined to be the union of the GO terms associated with each architecture in the set, 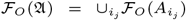. The functional similarity between a single architecture *A* and set of architectures *U* is

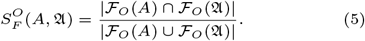

To avoid confusion due to use of the Jaccard similarity in two contexts, we use *functional similarity* to denote the similarity between two sets of GO codes and *Jaccard similarity* to refer to shared domain content.

### Annotated domain architecture neighborhoods

Given an embedding *E*, let *N*_*O,k*_(*A*) be the annotated nearest neighborhood of size *k* for architecture *A*, separately for each of the three sub-ontologies *O*. We define this *k-neighborhood* to be the *k* DAs in *A*_*O*_ that are closest to *A* in embedding *E*. Formally, 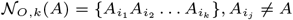 is a *k*-neighborhood of *A* if 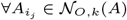

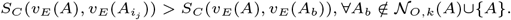

If there are two or more DAs that are equidistant from *A*, the k-neighborhood may not be unique. In this case, we assign one of the sets of *k* nearest neighbors to *N*_*O,k*_(*A*) arbitrarily.

We further consider the subsets of *N*_*O,k*_(*A*): the *shared k-neighborhood*, consisting of domain architectures that share at least one domain with *A*,

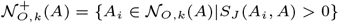

and the *non-sharing k-neighborhood*, consisting of domain architectures that share no domains with *A*,

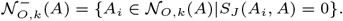

The respective sizes of these neighborhoods are *k*^+^ and *k*^−^, where *k*^+^ + *k*^−^ = *k*.

When comparing domain content with Jaccard similarity, without reference to an embedding, the *k*-neighborhood of *A* is defined analogously. In this case, all DAs in the *k*-neighborhood of *A* have at least one domain in common with *A*, and thus 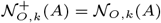 and 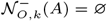.

### Functional similarity in the k-neighborhood

We ask whether multidomain architecture pairs that are close in the embedding tend to have similar functions. To assess whether domain architectures in the k-neighborhood of *A* have similar functions to *A*, we calculated 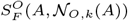, the functional similarity between *A* and its neighbors. The functional similarity is defined to be

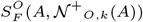

in shared neighborhoods and

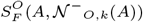

in non-sharing neighborhoods. For a given embedding and a given ontology, the mean functional similarity over all architectures in *A*_*O*_ for each *O* ∈ {MF, BP, CC} provides a measure of how well that embedding places domain architectures with similar functions in close proximity. We quantify performance using precision, recall, and Matthew’s Correlation Coefficients (MCC).

Functional consistency within shared k-neighborhoods.

To assess how tightly focused sets of nearest neighbors are, we computed two measures for each shared *k*=neighborhood. First, we considered the mean number of distinct GO terms in the shared *k*-neighborhood,

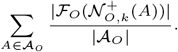

We also calculated 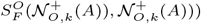, the mean functional similarity within the shared *k*-neighborhood obtained from all 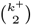 pairs of DAs in 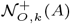,

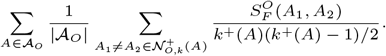

For each embedding method (or Jaccard), we report the mean value of each measure across all target domain architectures.

### Statistical tests for non-sharing nearest neighborhoods

We assessed the significance of observing neighboring domain architectures that are functionally similar, but have no common domains, using a randomization strategy with two test statistics.

Given an embedding and an ontology, let 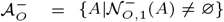 be the set of architectures in *A*_*O*_ that have at least one neighbor that with no common domain. The first test statistic is the mean functional similarity, averaged over all target architectures in 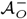,

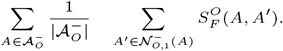

The second is the number of such pairs with functional similarity greater than 0.8:

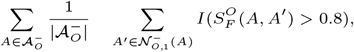

where *I* is the indicator function.

For each test statistic, the expected value was estimated from 100,000 null replicates generated as follows: for each target architecture in 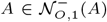, an non-sharing partner DA was selected at random, with replacement, from the set of all DAs that share no domain *A*. The value of the test statistic was calculated for set of null pairs generated in each replicate and averaged over all replicates to obtain the expected value. The empirical *p*-value is the proportion of replicates in which the null pairs achieve a greater value of the test statistic than the genuine pairs.

## Results

Here, we use vector semantics to investigate the relationship between protein domain content and protein function. We construct domain architecture embeddings using two sparse, high-dimensional encodings (TF-IDF, PMI) and six learned low-dimensional embeddings trained with Word2Vec. For each embedding, we ask: given a target domain architecture, how well do the combined GO terms of its *k* nearest annotated neighbors retrieve the GO annotations of the target? We assess this by measuring the functional similarity between the target and the union of GO terms from its neighbors. As shown in Table S1, functional similarity is high across all embeddings. In addition, GO term inheritance metrics - precision, recall, and MCC, are also high (Table S2). These results indicate that proximity in embedding space captures aspects of shared biological function.

To better understand the underlying functional signals captured by embeddings, we next consider two scenarios, when proximal domain architectures share at least one domain, and when they do not have any domain in common. These cases allow us to decompose the contribution of domain content and learned contextual patterns in domain organization.

### Nearest neighbors that share at least one domain

First, we examine the scenario where nearest neighbors share at least one domain with the target. This allows us to assess whether domain architecture embeddings provide more information than a simple domain content comparison. As a control, we calculated the same statistics using the *k* DAs with the highest Jaccard similarity to the target, that is, neighbors with the most similar domain content.

For multidomain architectures that share at least one domain, neighboring domain architectures have more similar GO annotations when the neighborhood is determined using a DA embedding than when domain content alone is considered (Fig. S1, Table S3). We compared how well GO terms in shared neighborhoods recapitulate the functional annotation of the target domain, for *k* = 1, 3, 5 and *k* = 10. For values of *k* greater than 1, The Word2Vec (*m* = 5, *w* = 5) embedding consistently obtained the best precision. The best recall was obtained when neighboring DAs were identified based on shared domain content using Jaccard similarity or TF-IDF, which also reflects shared domain content. This is also true when *k* = 1 for the MF and BP ontologies (Table 1).

**Table 1.** Accuracy of GO annotation transfer in the shared *k*-neighborhood for multidomain architectures as the mean 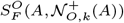 over all *A* ∈ *A*_*O*_.

| Molecular<br>Function | $k = 1$ | | | $k = 3$ | | | $k = 5$ | | | $k = 10$ | | |
| --- | --- | --- | --- | --- | --- | --- | --- | --- | --- | --- | --- | --- |
|  | precision | recall | MCC | precision | recall | MCC | precision | recall | MCC | precision | recall | MCC |
| TF-IDF | **0.829** | 0.816 | <u>0.806</u> | 0.671 | *0.884* | 0.745 | 0.580 | <u>0.907</u> | 0.695 | 0.455 | <u>0.932</u> | 0.615 |
| PMI | 0.789 | 0.796 | 0.778 | 0.651 | 0.874 | 0.732 | 0.573 | 0.898 | 0.69 | 0.471 | 0.921 | 0.625 |
| w2v(100,5) | 0.812 | 0.814 | 0.799 | 0.666 | <u>0.885</u> | *0.746* | 0.588 | *0.905* | 0.702 | 0.474 | 0.927 | 0.629 |
| w2v(100,1) | 0.792 | 0.793 | 0.778 | 0.649 | 0.869 | 0.728 | 0.570 | 0.894 | 0.686 | 0.475 | 0.917 | 0.626 |
| w2v(10,5) | <u>0.822</u> | *0.818* | <u>0.806</u> | *0.677* | 0.883 | <u>0.751</u> | *0.596* | 0.904 | <u>0.707</u> | *0.490* | *0.924* | *0.640* |
| w2v(10,1) | 0.809 | 0.804 | 0.792 | 0.669 | 0.877 | 0.743 | 0.588 | 0.896 | 0.699 | 0.488 | 0.918 | 0.636 |
| w2v(5,5) | 0.813 | <u>0.819</u> | 0.802 | **0.702** | 0.872 | **0.762** | **0.636** | 0.890 | **0.728** | **0.542** | 0.912 | **0.673** |
| w2v(5,1) | 0.792 | 0.793 | 0.777 | <u>0.679</u> | 0.853 | 0.739 | <u>0.618</u> | 0.872 | <u>0.707</u> | <u>0.540</u> | 0.898 | <u>0.665</u> |
| Jaccard | *0.819* | **0.823** | **0.808** | 0.643 | **0.902** | 0.736 | 0.530 | **0.926** | 0.668 | 0.382 | **0.944** | 0.559 |

| Biological<br>process | $k = 1$ | | | $k = 3$ | | | $k = 5$ | | | $k = 10$ | | |
| --- | --- | --- | --- | --- | --- | --- | --- | --- | --- | --- | --- | --- |
|  | precision | recall | MCC | precision | recall | MCC | precision | recall | MCC | precision | recall | MCC |
| TF-IDF | **0.799** | **0.794** | **0.780** | <u>0.644</u> | <u>0.859</u> | **0.719** | 0.546 | <u>0.883</u> | <u>0.662</u> | 0.415 | <u>0.910</u> | 0.576 |
| PMI | 0.741 | 0.747 | 0.727 | 0.591 | 0.832 | 0.673 | 0.513 | 0.863 | 0.631 | 0.413 | 0.891 | 0.565 |
| w2v(100,5) | 0.762 | 0.761 | 0.743 | 0.617 | 0.846 | 0.694 | 0.534 | *0.873* | 0.648 | 0.413 | *0.899* | 0.568 |
| w2v(100,1) | 0.741 | 0.740 | 0.723 | 0.605 | 0.837 | 0.683 | 0.526 | 0.861 | 0.638 | 0.422 | 0.889 | 0.572 |
| w2v(10,5) | *0.773* | *0.776* | *0.758* | 0.630 | *0.847* | *0.703* | *0.547* | *0.873* | *0.658* | 0.426 | 0.898 | 0.578 |
| w2v(10,1) | 0.757 | 0.759 | 0.741 | 0.619 | 0.842 | 0.694 | 0.545 | 0.870 | 0.655 | *0.430* | 0.895 | *0.579* |
| w2v(5,5) | 0.768 | 0.765 | 0.750 | **0.653** | 0.825 | <u>0.709</u> | **0.589** | 0.854 | **0.679** | **0.486** | 0.879 | **0.616** |
| w2v(5,1) | 0.755 | 0.759 | 0.738 | *0.634* | 0.817 | 0.691 | <u>0.570</u> | 0.833 | 0.656 | <u>0.481</u> | 0.869 | <u>0.607</u> |
| Jaccard | <u>0.792</u> | <u>0.787</u> | <u>0.772</u> | 0.604 | **0.871** | 0.698 | 0.487 | **0.896** | 0.626 | 0.310 | **0.922** | 0.494 |

| Cellular<br>component | $k = 1$ | | | $k = 3$ | | | $k = 5$ | | | $k = 10$ | | |
| --- | --- | --- | --- | --- | --- | --- | --- | --- | --- | --- | --- | --- |
|  | precision | recall | MCC | precision | recall | MCC | precision | recall | MCC | precision | recall | MCC |
| TF-IDF | *0.891* | <u>0.886</u> | <u>0.874</u> | 0.782 | **0.926** | <u>0.829</u> | *0.710* | **0.939** | *0.790* | *0.611* | <u>0.953</u> | *0.727* |
| PMI | 0.871 | 0.856 | 0.848 | 0.756 | 0.911 | 0.805 | 0.687 | 0.925 | 0.766 | 0.598 | 0.941 | 0.712 |
| w2v(100,5) | 0.886 | 0.879 | 0.868 | 0.756 | 0.922 | 0.808 | 0.695 | *0.937* | 0.776 | 0.593 | 0.948 | 0.710 |
| w2v(100,1) | 0.877 | 0.868 | 0.858 | 0.761 | 0.921 | 0.813 | 0.698 | 0.934 | 0.776 | *0.611* | 0.946 | 0.722 |
| w2v(10,5) | 0.888 | <u>0.886</u> | <u>0.874</u> | *0.785* | <u>0.924</u> | <u>0.829</u> | 0.703 | *0.937* | 0.782 | 0.603 | *0.952* | 0.720 |
| w2v(10,1) | 0.874 | 0.885 | 0.865 | 0.760 | 0.920 | 0.811 | 0.701 | 0.934 | 0.778 | 0.608 | 0.949 | 0.722 |
| w2v(5,5) | **0.895** | **0.890** | **0.879** | **0.808** | 0.921 | **0.841** | **0.742** | 0.932 | **0.804** | **0.651** | 0.941 | **0.747** |
| w2v(5,1) | 0.886 | 0.883 | 0.870 | <u>0.786</u> | 0.910 | 0.823 | <u>0.732</u> | 0.923 | <u>0.794</u> | <u>0.648</u> | 0.937 | <u>0.744</u> |
| Jaccard | <u>0.893</u> | 0.869 | 0.866 | 0.752 | <u>0.924</u> | 0.810 | 0.646 | <u>0.938</u> | 0.747 | 0.484 | **0.955** | 0.638 |

Best-performing method for each metric in bold, second-best is underlined, and third-best is italicized.

This led us to hypothesize that shared domain content provides some functional information, but is not sufficiently specific. That is, some DAs that share domains have similar functions, but others do not. To investigate this, we compared the number of distinct GO terms in the shared-domain neighborhood (*k* = 3 and *k* = 5) obtained with Jaccard and with the various embeddings. The number of terms retrieved using Jaccard similarity exceeds the number of terms retrieved using all embeddings in all three ontologies, and is larger than the best embedding by as much as 50% (Table 2). Similarly, mean GO term similarity between domain architecture pairs within the shared-domain neighborhood is consistently smaller with Jaccard, compared with all other methods (Table S4). These results suggest that shared domain content alone is not sufficiently precise. Embeddings are deriving more information than simply the presence or absence of shared domains.

**Table 2.** functional consistency within shared *k*-neighborhoods measured as the mean 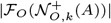 over all *A* ∈ *A*_*O*_.

| Embedding | MF |  | BP |  | CC |  |
| --- | --- | --- | --- | --- | --- | --- |
| | $k = 3$ | $k = 5$ | $k = 3$ | $k = 5$ | $k = 3$ | $k = 5$ |
| TF-IDF | 15.0 | 18.6 | 21.5 | 27.6 | 8.8 | 10.3 |
| PMI | 15.2 | 18.4 | 21.4 | 27.3 | 8.5 | 10.1 |
| w2v(100,5) | 15.1 | 18.4 | 21.9 | 28.0 | 9.3 | 10.9 |
| w2v(100,1) | 15.2 | 18.8 | 21.7 | 27.5 | 8.9 | 10.6 |
| w2v(10,5) | 14.6 | 18.0 | 21.2 | 27.2 | 8.7 | 10.5 |
| w2v(10,1) | 14.7 | 18.0 | 21.3 | 26.9 | 9.0 | 10.6 |
| w2v(5,5) | 12.5 | 15.3 | 16.7 | 21.0 | 7.4 | 8.9 |
| w2v(5,1) | 12.2 | 15.1 | 16.8 | 21.3 | 7.3 | 8.8 |
| Jaccard | 17.1 | 22.7 | 25.0 | 34.0 | 10.0 | 12.5 |

### Nearest neighbors that share no domains

Embedding-based document similarity can identify texts on similar subjects even when they do not contain the same words. This is because they can contain words with similar or related meanings, captured by the embedding. By analogy, we ask whether embeddings are able to identify domain architectures with similar functions, even when they share no domains.

To explore this, we first asked how often multidomain architectures that share no domain are nearest neighbors in an embedding. Indeed, as many as 30% of proximal multidomain architectures have no domains in common (Table 3), depending on the embedding and the ontology. TF-IDF retrieves notably fewer such architectures than other embeddings, which is not surprising given that TF-IDF explicitly considers domains that are shared across architectures.

**Table 3.**
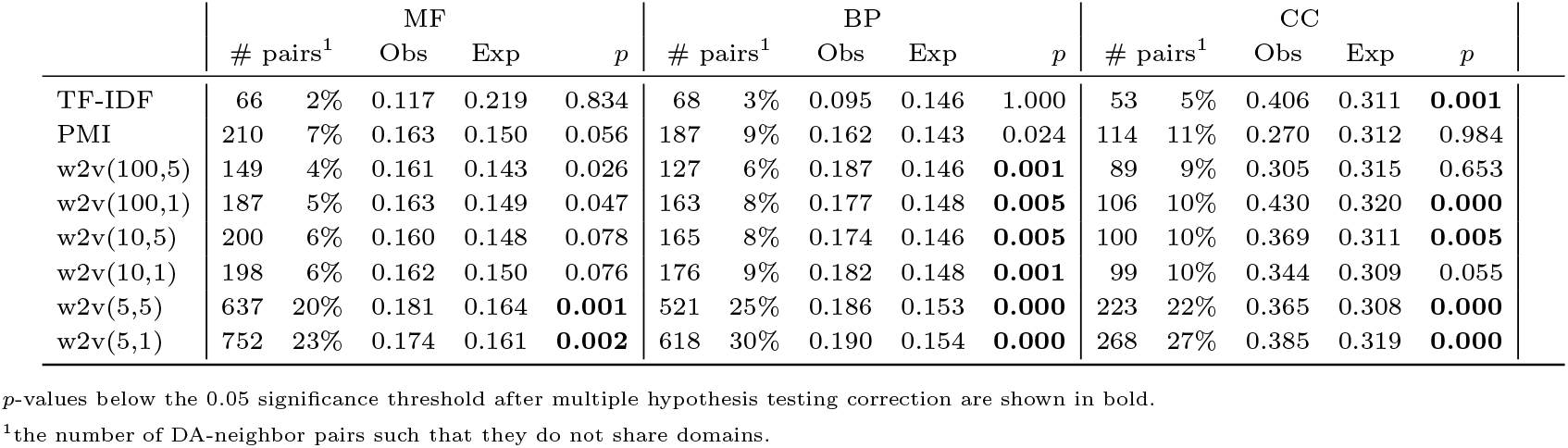
Mean functional similarity 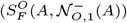 averaged over all *A* ∈ *A*_*O*_) in non-sharing neighborhoods (*k* = 1).

We next asked whether proximal domain architectures that share no domain with the target have similar GO terms (Fig. 1). For BP and CC, the majority of embeddings (shown in bold in Table 3) achieve significantly higher mean functional similarity than expected by chance. For MF, two embeddings identify non-sharing neighbors with higher than expected similarity. The observed mean similarity exceeds that of the null model for all embeddings except TF-IDF. Larger neighborhoods (*k* = 3, 5, 10) tell a similar story: non-sharing neighbors remain highly functionally relevant (Fig. S2, Table S5).

To further assess whether embeddings identify non-sharing neighbors with meaningful similarities, we asked how often such pairs have high functional similarity. Indeed, embeddings consistently identify more pairs with functional similarity greater than 0.8 than expected by chance (Table 4). The results are statistically significant for w2v(5,1) in all three ontologies,for w2v(5,5) in BP and CC, and additionally w2v(100,1) and w2v(10,5) for CC.

**Table 4.** Number of neighboring multidomain pairs (*k* = 1) that lack a common domain with functional similarity *>* 0.8.

|  | MF |  |  | BP |  |  |
| --- | --- | --- | --- | --- | --- | --- |
| | Obs | Exp | $p$ | Obs | Exp | $p$ |
| TF-IDF | 0 | 0.1 | 0.134 | 0 | 0.3 | 0.298 |
| PMI | 2 | 0.9 | 0.062 | 6 | 1.2 | **0.000** |
| w2v(100,5) | 2 | 0.6 | 0.018 | 2 | 0.9 | 0.065 |
| w2v(100,1) | 3 | 0.8 | 0.007 | 4 | 1.5 | 0.018 |
| w2v(10,5) | 0 | 0.7 | 0.522 | 2 | 1.3 | 0.148 |
| w2v(10,1) | 0 | 0.7 | 0.519 | 4 | 1.4 | 0.012 |
| w2v(5,5) | 9 | 4.1 | 0.008 | 15 | 5.4 | **0.000** |
| w2v(5,1) | 11 | 4.4 | **0.002** | 15 | 6.7 | **0.001** |

|  | CC |  |  |
| --- | --- | --- | --- |
| | Obs | Exp | $p$ |
| TF-IDF | 8 | 4.1 | 0.010 |
| PMI | 5 | 9.0 | 0.912 |
| w2v(100,5) | 8 | 7.7 | 0.356 |
| w2v(100,1) | 19 | 9.1 | **0.000** |
| w2v(10,5) | 15 | 7.8 | **0.002** |
| w2v(10,1) | 10 | 7.3 | 0.096 |
| w2v(5,5) | 26 | 16.4 | **0.004** |
| w2v(5,1) | 42 | 22.0 | **0.000** |

$p$ -values below the 0.05 significance threshold after multiple hypothesis testing correction are shown in bold.

The possibility that domain architecture embeddings can identify multidomain pairs that share functional annotations, even when there is no overlapping domain content, is intriguing. To further explore this result, we examined the 187 unique pairs of nearest neighbors (*k* = 1) that lack a shared domain, but have functional similarity above 0.8, identified by any embedding. Interestingly, only a small number of these 187 pairs are found by more than one embedding. This low overlap suggests that each embedding captures different contextual signals, highlighting complementary aspects of functional relationships between domain architectures. The complete list of such pairs is provided in Supplementary Information (S1). The threshold of 0.8 was selected to be consistent with the mean functional similarity observed in nearest neighbors (*k* = 1) that share at least one domain (Table S3). The best threshold for identifying meaningful similarity is a question for future work.

DA pairs that lack shared domains, but nevertheless have high functional similarity, might be caused by errors in domain annotation. To screen for such cases, we compared the amino acid sequences that correspond to the 187 pairs with functional similarity greater than 0.8, in search of conserved regions that might correspond to a shared domain that was overlooked. Out of 14,486 pairwise sequence comparisons (recall that most DAs correspond to more than one sequence) only 16 sequence pairs had local alignments with an E value less than 1. Only two domain architecture pairs corresponded to sequence comparisons with E values below 1 (E = 0.77 and E= 0.074, respectively). At time of writing, the default significance threshold in the blastp interface is (E *<* 0.05).

We discuss two of these pairs in greater detail. The Zinc finger transcription factor Trps1 and Homeobox Hox-B4 proteins (Figure 2, top panel) are both transcription factors involved in developmental regulation of the skeletal system [Kaiser et al., 2007, Dias et al., 2013]. Trps1 is associated with regulation of chondrocyte differentiation (GO:0032330), which is “part of” skeletal system development (GO:0001501). Mutations in the gene are the basis of tricho-rhino-phalangeal syndrome type I (TRPS I) a genetic disorder characterized by skeletal abnormalities [Momeni et al., 2000]. *HOXB4* is associated with embryonic skeletal system morphogenesis (GO:0048704) and bone marrow development (GO:0048539), also “part of” skeletal system development. *HOXB4* plays crucial roles in vertebrate skeletal system development (reviewed in Morgan et al. [2004]) and regulates the balance between differentiation into osteogenic (bone formation) or hematopoietic lineages of human embryonic stem cells [Kärner et al., 2009]. As transcription factors, *TRPS1* and *HOXB4* also share GO terms such as regulation of RNA biosynthetic process (GO:2001141). Both domain architectures contain DNA-binding domains (glucocorticoid receptor-like domains, zinc fingers, and homeodomains), but not the same DNA-binding domains. The blastp web interface identifies no significant similarity between these sequences at default settings.

**Fig. 2.**
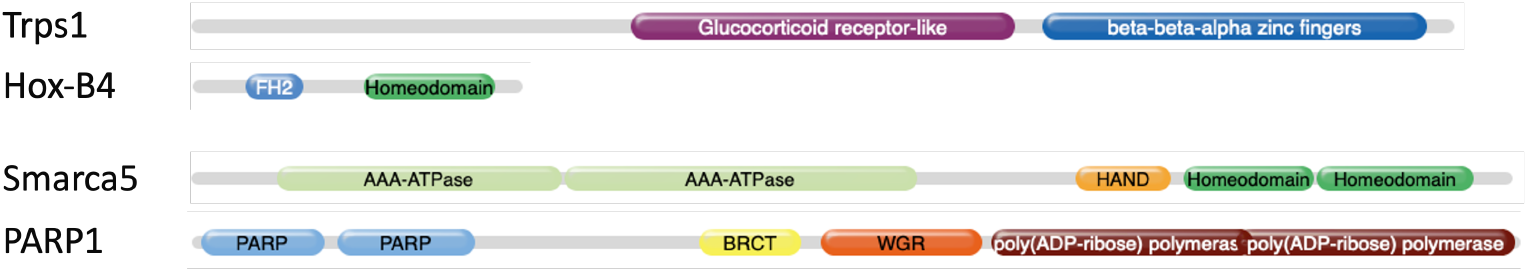
Schematic representation of domain architectures of two pairs of multidomain proteins that share no domains, but have high functional term similarity (see text). The relative lengths of the proteins and domains are approximate.

SWI/SNF-related matrix-associated actin-dependent regulator of chromatin subfamily A member 5 (Smarca5) and Poly [ADP-ribose] polymerase (PARP) (Figure 2, bottom panel) are DNA-binding proteins that participate in DNA repair [Xie et al., 2015, Toiber et al., 2013]. They share CC GO terms (nucleolus (GO:0005730) and site of double-strand break (GO:0035861), and MF GO terms (DNA binding (GO:0003677)). Both domain architectures encode DNA binding domains (HAND domains and homeodomains versus PARP zinc fingers and WGR domains), as well as domains with enzymatic activities (AAA-ATPases and poly(ADP-ribose)polymerases, respectively), but the specific domains in each of these categories differ in the two proteins. This pair has no significant sequence similarity at default parameter settings.

In this pilot study, vector embeddings identified neighboring domain architecture pairs that share functional properties, but lack a common domain. There are more such pairs than expected by chance and the mean similarity of nearest neighbors that lack a shared domain is also greater than expected. However, many non-sharing nearest neighbors do not have strong functional similarity. Additional research is needed to determine how to identify the most promising pairs.

## Discussion

Mounting evidence suggests that protein domains carry information about the functions of the proteins that encode them, but explicit models that relate domain content to protein function are lacking. In particular, we require a better understanding of how the functional relevance of a particular domain depends on its surrounding context. Here, we investigated the potential of vector semantic embeddings to capture contextual information. We observe that pairs of multidomain architectures identified using vector semantics share more functional attributes than pairs of multidomain architectures selected based on domain content similarity, alone. Our results suggest that these vector semantic models may capture combinatorial interactions of domains in the same protein. Surprisingly, in some cases, vector semantic embeddings identified multidomain architecture pairs that had high functional similarity, *despite having no domains in common at all*. Note that no information from sequence analysis or biological textual descriptions were used to construct the embeddings; these inferences are based on domain co-occurrence alone. Taken together, these results underscore the potential of protein domain vector semantics to elucidate the “design rules” of multidomain architectures.

While intriguing, these results are preliminary. Further investigations that comprehend larger data sets and greater taxonomic breadth are required. The revolution in protein structure prediction driven by advances in deep learning has produced greatly expanded bioinformatic resources for protein domain analyses [Paysan-Lafosse et al., 2025, Lau et al., 2024]. Future work that incorporates this new knowledge is also imperative.

The results presented here suggest that protein domain vector semantics may be able to discover the protein domain equivalent of words and phrases with similar or related meanings in natural languages. The science of lexical semantics deals with many types of related meanings [reviewed in Jurafsky and Martin, 2008]. Words may have similar meanings (e.g., *sarcastic, ironic*) be used in the same context (e.g., *coffee, cup*), be made of the same materials or have the similar constituent parts (e.g., *car, bicycle*), or be associated with the same activity (e.g., *suture, scalpel*). Characterizing different types of word associations and developing NLP models that are capable of distinguishing between them are active areas of NLP research [Hill et al., 2015]. Our discovery of functionally similar multidomain architectures that harbor no common domains suggests that notions of semantic or topical relatedness may be relevant to protein domain function. What types of “topical relatedness” are meaningful in the protein domain context is an exciting open question. In one of the first works to tackle this question, Buchan and Jones [2020] experimented with vector arithmetic models of word analogies (e.g., *king* is to *man* as *queen* is to *woman*) to explore whether such relations also exist for domains.

Vector semantic models of multidomain proteins also hold promise for advancing bioinformatic applications. The problem of developing a *Domain Ontology* is closely related to questions in protein domain lexical semantics. The Gene Ontology (GO) is a powerful and general system for describing protein function. The hierarchy of well-defined functional terms using a controlled vocabulary supports a broad range of applications to query, manipulate, compare and reason with biological data. Currently, no equivalent domain function ontology exists; nor is there agreement on what a domain-specific ontology should look like. Should domain ontology terms describe inherent functional properties of the domains themselves? Or is the goal to annotate domains with terms that predict the functions of any protein that is found to encode the domain? Several lines of research are advancing functional annotation for domains. The InterPro2GO project is building a manually annotated domain resource, where annotations are based on experimental evidence of a domain’s function, and not simply the function of its associated proteins [Burge et al., 2012, Blum et al., 2021]. In an orthogonal approach, strategies are being developed to support automated mapping of Gene Ontology (GO) terms from proteins to domains [Weiner et al., 2008, Buchan and Jones, 2020, Fang and Gough, 2013, López and Pazos, 2013, Ulusoy and Doğan, 2024, Ibtehaz et al., 2023]. However, these annotations are couched in terms of the GO, which was developed to describe the functions of genes and proteins, not domains. Further examination of the appropriateness of the MF, BP, and CC trichotomy for describing domain function is warranted. It may be possible to partition the molecular functions of a protein by attributing specific molecular roles to domains, but it is less clear whether protein domains tend to be associated with specific biological processes or cellular compartments. This is an interesting research question that has practical consequences for the development of bioinformatic resources.

Vector semantics of domains and domain architectures also holds promise for *protein function prediction* [Radivojac et al., 2013, Jiang et al., 2016, Zhou et al., 2019]. A better understanding of multidomain design rules will shed light on the predictive value of domain content and organization for multidomain protein functions. This is important: In the first Critical Assessment of protein Function Annotation (CAFA) challenge, all of the top predictors performed less well on multidomain proteins than on single domain proteins [Radivojac et al., 2013]. Protein function prediction methods are increasingly combining many different types of data [Radivojac et al., 2013, Jiang et al., 2016, Zhou et al., 2019, Boadu et al., 2025], including domain family information, but few methods have attempted to incorporate an explicit model relating domain content to the functional attributes of multidomain proteins. The success of several notable exceptions [You et al., 2018, Ibtehaz et al., 2023, Talo and Bozdag, 2025] suggests the value of vector semantics in this context.

## Data availability

The data and code used in this work are available on Zenodo (draft).

