## Supplementary material for "Vector Semantics of Multidomain Protein Architectures": GLBIO2025_CuiXiaoStolzerDurand_supp.pdf

### Supplementary information for Vector Semantics of Multidomain Protein Architectures

**Table S1.** Mean functional similarity between nearest neighbors, ($S_F^O(\mathcal{N}_{O,k}(A), \mathcal{N}_{O,k}(A))$ averaged over all $A \in \mathcal{A}_O$), regardless of domain content sharing.

| MF | $k=1$ | $k=3$ | $k=5$ | $k=10$ |
|---|---|---|---|---|
| TF-IDF | 0.725 | 0.645 | 0.597 | 0.530 |
| PMI | 0.654 | 0.574 | 0.531 | 0.476 |
| w2v(100,5) | 0.690 | 0.611 | 0.569 | 0.511 |
| w2v(100,1) | 0.667 | 0.582 | 0.538 | 0.482 |
| w2v(10,5) | 0.681 | 0.610 | 0.569 | 0.513 |
| w2v(10,1) | 0.670 | 0.594 | 0.553 | 0.501 |
| w2v(5,5) | 0.589 | 0.519 | 0.486 | 0.443 |
| w2v(5,1) | 0.558 | 0.493 | 0.466 | 0.427 |

| BP | $k=1$ | $k=3$ | $k=5$ | $k=10$ |
|---|---|---|---|---|
| TF-IDF | 0.679 | 0.578 | 0.522 | 0.446 |
| PMI | 0.573 | 0.486 | 0.448 | 0.396 |
| w2v(100,5) | 0.616 | 0.528 | 0.487 | 0.430 |
| w2v(100,1) | 0.594 | 0.502 | 0.458 | 0.405 |
| w2v(10,5) | 0.617 | 0.538 | 0.493 | 0.434 |
| w2v(10,1) | 0.604 | 0.516 | 0.479 | 0.422 |
| w2v(5,5) | 0.523 | 0.461 | 0.430 | 0.388 |
| w2v(5,1) | 0.498 | 0.431 | 0.404 | 0.366 |

| CC | $k=1$ | $k=3$ | $k=5$ | $k=10$ |
|---|---|---|---|---|
| TF-IDF | 0.775 | 0.694 | 0.652 | 0.597 |
| PMI | 0.692 | 0.621 | 0.586 | 0.538 |
| w2v(100,5) | 0.739 | 0.656 | 0.623 | 0.576 |
| w2v(100,1) | 0.729 | 0.641 | 0.603 | 0.556 |
| w2v(10,5) | 0.739 | 0.664 | 0.629 | 0.580 |
| w2v(10,1) | 0.725 | 0.641 | 0.611 | 0.565 |
| w2v(5,5) | 0.684 | 0.611 | 0.583 | 0.544 |
| w2v(5,1) | 0.655 | 0.577 | 0.554 | 0.524 |

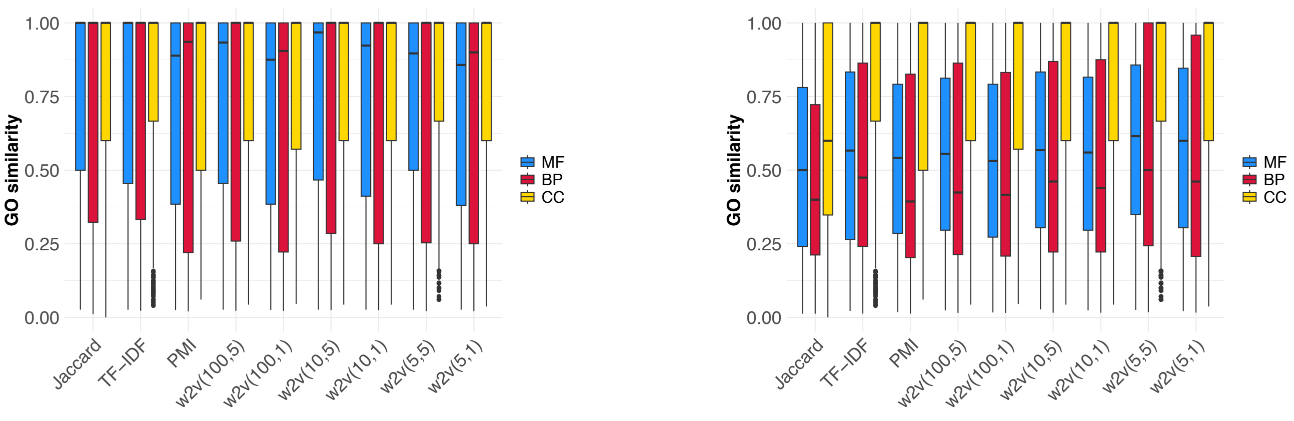


**Fig. S1.** Functional similarity between target domain architectures and the nearest neighbors when $k=1$ (left) or $k=5$ (right) that share at least one domain. GO terms associated with each target are compared to the union of GO terms from neighbors.

**Table S2.** Accuracy of GO annotation transfer in the $k$-neighborhood for multidomain architectures as the mean $S_F^O(A, \mathcal{N}_{O,k}(A))$ over all $A \in \mathcal{A}_O$, regardless of domain content sharing.

| **Molecular Function** | $k=1$ | | | $k=3$ | | | $k=5$ | | | $k=10$ | | |
|---|---|---|---|---|---|---|---|---|---|---|---|---|
| | precision | recall | MCC | precision | recall | MCC | precision | recall | MCC | precision | recall | MCC |
| TF-IDF | 0.827 | 0.811 | 0.796 | 0.671 | 0.879 | 0.735 | 0.568 | 0.902 | 0.675 | 0.410 | 0.926 | 0.572 |
| PMI | 0.757 | 0.761 | 0.735 | 0.569 | 0.844 | 0.659 | 0.463 | 0.874 | 0.596 | 0.331 | 0.905 | 0.499 |
| w2v(100,5) | 0.787 | 0.789 | 0.766 | 0.615 | 0.861 | 0.697 | 0.511 | 0.884 | 0.635 | 0.374 | 0.910 | 0.541 |
| w2v(100,1) | 0.772 | 0.767 | 0.746 | 0.582 | 0.844 | 0.668 | 0.471 | 0.872 | 0.602 | 0.343 | 0.903 | 0.512 |
| w2v(10,5) | 0.783 | 0.778 | 0.758 | 0.619 | 0.858 | 0.698 | 0.517 | 0.882 | 0.638 | 0.385 | 0.908 | 0.548 |
| w2v(10,1) | 0.776 | 0.767 | 0.748 | 0.598 | 0.846 | 0.678 | 0.497 | 0.873 | 0.620 | 0.368 | 0.902 | 0.532 |
| w2v(5,5) | 0.705 | 0.704 | 0.677 | 0.529 | 0.809 | 0.618 | 0.441 | 0.846 | 0.571 | 0.325 | 0.888 | 0.493 |
| w2v(5,1) | 0.680 | 0.676 | 0.649 | 0.499 | 0.786 | 0.590 | 0.411 | 0.831 | 0.544 | 0.304 | 0.878 | 0.472 |

| **Biological process** | $k=1$ | | | $k=3$ | | | $k=5$ | | | $k=10$ | | |
|---|---|---|---|---|---|---|---|---|---|---|---|---|
| | precision | recall | MCC | precision | recall | MCC | precision | recall | MCC | precision | recall | MCC |
| TF-IDF | 0.762 | 0.770 | 0.741 | 0.590 | 0.845 | 0.674 | 0.490 | 0.864 | 0.614 | 0.343 | 0.902 | 0.515 |
| PMI | 0.657 | 0.679 | 0.644 | 0.480 | 0.777 | 0.577 | 0.391 | 0.819 | 0.526 | 0.272 | 0.864 | 0.441 |
| w2v(100,5) | 0.709 | 0.710 | 0.687 | 0.532 | 0.802 | 0.622 | 0.437 | 0.836 | 0.568 | 0.309 | 0.874 | 0.479 |
| w2v(100,1) | 0.690 | 0.690 | 0.667 | 0.503 | 0.792 | 0.599 | 0.403 | 0.823 | 0.539 | 0.284 | 0.865 | 0.455 |
| w2v(10,5) | 0.710 | 0.708 | 0.686 | 0.544 | 0.805 | 0.630 | 0.446 | 0.836 | 0.574 | 0.316 | 0.873 | 0.484 |
| w2v(10,1) | 0.700 | 0.694 | 0.675 | 0.521 | 0.790 | 0.610 | 0.436 | 0.828 | 0.563 | 0.305 | 0.867 | 0.473 |
| w2v(5,5) | 0.627 | 0.628 | 0.602 | 0.471 | 0.753 | 0.563 | 0.392 | 0.800 | 0.523 | 0.278 | 0.853 | 0.447 |
| w2v(5,1) | 0.603 | 0.612 | 0.582 | 0.436 | 0.734 | 0.533 | 0.360 | 0.782 | 0.493 | 0.257 | 0.840 | 0.426 |

| **Cellular component** | $k=1$ | | | $k=3$ | | | $k=5$ | | | $k=10$ | | |
|---|---|---|---|---|---|---|---|---|---|---|---|---|
| | precision | recall | MCC | precision | recall | MCC | precision | recall | MCC | precision | recall | MCC |
| TF-IDF | 0.842 | 0.851 | 0.829 | 0.696 | 0.909 | 0.764 | 0.596 | 0.925 | 0.702 | 0.464 | 0.944 | 0.609 |
| PMI | 0.781 | 0.791 | 0.762 | 0.622 | 0.875 | 0.702 | 0.518 | 0.898 | 0.636 | 0.386 | 0.929 | 0.542 |
| w2v(100,5) | 0.828 | 0.819 | 0.803 | 0.661 | 0.881 | 0.729 | 0.569 | 0.901 | 0.675 | 0.434 | 0.928 | 0.585 |
| w2v(100,1) | 0.824 | 0.806 | 0.794 | 0.649 | 0.879 | 0.721 | 0.554 | 0.898 | 0.664 | 0.416 | 0.922 | 0.570 |
| w2v(10,5) | 0.829 | 0.815 | 0.802 | 0.684 | 0.878 | 0.742 | 0.592 | 0.900 | 0.690 | 0.445 | 0.927 | 0.592 |
| w2v(10,1) | 0.812 | 0.807 | 0.789 | 0.658 | 0.874 | 0.725 | 0.568 | 0.900 | 0.674 | 0.431 | 0.928 | 0.582 |
| w2v(5,5) | 0.783 | 0.777 | 0.757 | 0.625 | 0.863 | 0.698 | 0.530 | 0.890 | 0.644 | 0.393 | 0.923 | 0.551 |
| w2v(5,1) | 0.766 | 0.752 | 0.733 | 0.578 | 0.846 | 0.661 | 0.497 | 0.884 | 0.620 | 0.380 | 0.916 | 0.540 |

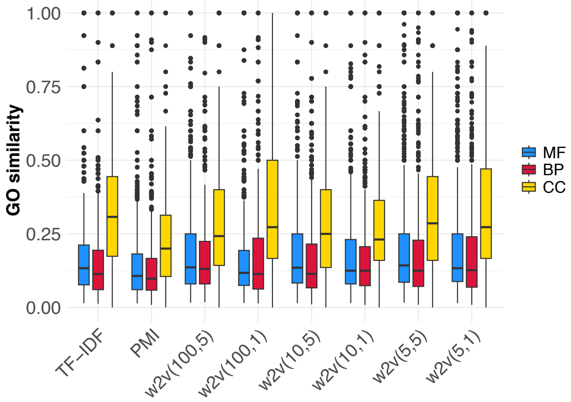


**Fig. S2.** Functional similarity between target domain architectures and the $k=5$ nearest neighbor that lacks a common domain. GO terms associated with each target are compared to the union of GO terms from neighbors.

#### DA pairs that lack shared domains but have high functional similarity

The list of pairs of DAs that are identified as nearest neighbors ($k=1$) that lack a shared domain, but have functional, similarity above 0.8, is provided as candidates.xlsx at `https://zenodo.org/records/15769961/preview/zenodo.zip?preview=1&include_deleted=0#tree_item5`

**Table S3.** Mean functional similarity within shared $k$-neighborhoods, $(S_F^O(\mathcal{N}_{O,k}^+(A), \mathcal{N}_{O,k}^+(A))$ averaged over all $A \in \mathcal{A}_O)$.

| MF | $k=1$ | $k=3$ | $k=5$ | $k=10$ |
|---|---|---|---|---|
| TF-IDF | 0.753 | 0.657 | 0.576 | 0.436 |
| PMI | 0.717 | 0.622 | 0.548 | 0.442 |
| w2v(100,5) | 0.746 | 0.649 | 0.560 | 0.447 |
| w2v(100,1) | 0.724 | 0.624 | 0.542 | 0.439 |
| w2v(10,5) | 0.752 | 0.661 | 0.574 | 0.466 |
| w2v(10,1) | 0.736 | 0.642 | 0.566 | 0.457 |
| w2v(5,5) | 0.740 | 0.659 | 0.601 | 0.507 |
| w2v(5,1) | 0.725 | 0.642 | 0.580 | 0.497 |
| Jaccard | 0.754 | 0.637 | 0.524 | 0.375 |

| BP | $k=1$ | $k=3$ | $k=5$ | $k=10$ |
|---|---|---|---|---|
| TF-IDF | 0.724 | 0.613 | 0.526 | 0.403 |
| PMI | 0.662 | 0.559 | 0.492 | 0.399 |
| w2v(100,5) | 0.680 | 0.583 | 0.511 | 0.399 |
| w2v(100,1) | 0.657 | 0.570 | 0.501 | 0.406 |
| w2v(10,5) | 0.699 | 0.595 | 0.524 | 0.411 |
| w2v(10,1) | 0.678 | 0.585 | 0.522 | 0.415 |
| w2v(5,5) | 0.689 | 0.608 | 0.557 | 0.464 |
| w2v(5,1) | 0.672 | 0.588 | 0.534 | 0.459 |
| Jaccard | 0.717 | 0.584 | 0.476 | 0.305 |

| CC | $k=1$ | $k=3$ | $k=5$ | $k=10$ |
|---|---|---|---|---|
| TF-IDF | 0.827 | 0.749 | 0.686 | 0.596 |
| PMI | 0.794 | 0.718 | 0.659 | 0.579 |
| w2v(100,5) | 0.820 | 0.719 | 0.668 | 0.573 |
| w2v(100,1) | 0.807 | 0.725 | 0.670 | 0.590 |
| w2v(10,5) | 0.826 | 0.750 | 0.677 | 0.586 |
| w2v(10,1) | 0.816 | 0.724 | 0.674 | 0.589 |
| w2v(5,5) | 0.833 | 0.767 | 0.710 | 0.623 |
| w2v(5,1) | 0.821 | 0.742 | 0.695 | 0.621 |
| Jaccard | 0.817 | 0.722 | 0.628 | 0.476 |

**Table S4.** Average pairwise GO similarity between neighbors in neighborhood of size $k$, excluding the target domain architecture.

| | MF | | BP | | CC | |
|---|---|---|---|---|---|---|
| Embedding | $k=3$ | $k=5$ | $k=3$ | $k=5$ | $k=3$ | $k=5$ |
| TF-IDF | 0.68 | 0.64 | 0.61 | 0.56 | 0.76 | 0.73 |
| PMI | 0.65 | 0.61 | 0.57 | 0.54 | 0.75 | 0.72 |
| w2v(100,5) | 0.67 | 0.63 | 0.59 | 0.55 | 0.74 | 0.72 |
| w2v(100,1) | 0.64 | 0.61 | 0.57 | 0.54 | 0.73 | 0.72 |
| w2v(10,5) | 0.69 | 0.64 | 0.61 | 0.57 | 0.77 | 0.74 |
| w2v(10,1) | 0.67 | 0.64 | 0.59 | 0.56 | 0.74 | 0.72 |
| w2v(5,5) | 0.69 | 0.66 | 0.65 | 0.61 | 0.78 | 0.75 |
| w2v(5,1) | 0.67 | 0.64 | 0.61 | 0.59 | 0.74 | 0.73 |
| Jaccard | 0.61 | 0.57 | 0.54 | 0.49 | 0.70 | 0.67 |

**Table S5.** Average functional similarity between nearest neighbors that lack a common domain.

| MF | $k = 1$ | $k = 3$ | $k = 5$ | $k = 10$ |
|---|---|---|---|---|
| TF-IDF | 0.114 | 0.188 | 0.188 | 0.192 |
| PMI | 0.161 | 0.153 | 0.153 | 0.147 |
| w2v(100,5) | 0.178 | 0.194 | 0.215 | 0.199 |
| w2v(100,1) | 0.184 | 0.179 | 0.172 | 0.168 |
| w2v(10,5) | 0.169 | 0.191 | 0.203 | 0.205 |
| w2v(10,1) | 0.188 | 0.183 | 0.196 | 0.178 |
| w2v(5,5) | 0.214 | 0.209 | 0.207 | 0.188 |
| w2v(5,1) | 0.213 | 0.220 | 0.207 | 0.187 |

| BP | $k = 1$ | $k = 3$ | $k = 5$ | $k = 10$ |
|---|---|---|---|---|
| TF-IDF | 0.095 | 0.168 | 0.168 | 0.174 |
| PMI | 0.162 | 0.140 | 0.143 | 0.126 |
| w2v(100,5) | 0.187 | 0.173 | 0.179 | 0.174 |
| w2v(100,1) | 0.177 | 0.180 | 0.176 | 0.166 |
| w2v(10,5) | 0.174 | 0.180 | 0.182 | 0.170 |
| w2v(10,1) | 0.182 | 0.170 | 0.170 | 0.163 |
| w2v(5,5) | 0.186 | 0.182 | 0.181 | 0.180 |
| w2v(5,1) | 0.190 | 0.188 | 0.181 | 0.174 |

| CC | $k = 1$ | $k = 3$ | $k = 5$ | $k = 10$ |
|---|---|---|---|---|
| TF-IDF | 0.406 | 0.381 | 0.346 | 0.355 |
| PMI | 0.270 | 0.265 | 0.237 | 0.216 |
| w2v(100,5) | 0.305 | 0.307 | 0.314 | 0.311 |
| w2v(100,1) | 0.430 | 0.342 | 0.355 | 0.320 |
| w2v(10,5) | 0.369 | 0.334 | 0.331 | 0.301 |
| w2v(10,1) | 0.344 | 0.307 | 0.303 | 0.288 |
| w2v(5,5) | 0.365 | 0.350 | 0.341 | 0.290 |
| w2v(5,1) | 0.385 | 0.354 | 0.341 | 0.308 |
